# Adaptive Sampling for Spatial Capture-Recapture: An efficient sampling scheme for rare or patchily distributed species

**DOI:** 10.1101/357459

**Authors:** Alec Wong, Angela Fuller, J. Andrew Royle

## Abstract

Rare species present challenges to data collection, particularly when the species is spatially clustered over large areas, such that the encounter frequency of the organism is low. Sampling where the organism is absent consumes resources, and offers relatively low-quality information which are often difficult to model using standard statistical methods. In adaptive sampling, a probabilistic sampling method is employed first, and additional effort is allocated in the vicinity of sites where some measured index variable - assumed to be proportional to local population size - exceeds an *a priori* threshold. We applied this principle to the spatial capture-recapture (SCR) analytical framework in a Bayesian hierarchical model incorporating capture-recapture (CR) and index information from unsampled sites to estimate density. We assessed the adaptively sampled SCR model (AS-SCR) by simulating CR data and compared performance with a standard SCR baseline (F-SCR), adaptive SCR discarding index information (AS-SCR–), and standard SCR applied at a simple random sample of sites. Under AS-SCR, we observed minimal bias and comparable variance with respect to parameter estimates provided by the standard F-SCR model and sampling implementation, but with substantially reduced effort and significant cost saving potential. This represents the first application of adaptive sampling to SCR.

## Introduction

Knowledge of species abundance is a critical component of wildlife conservation and management (Williams, Nichols, and Conroy 2002), and monitoring this state variable is especially important when managing rare or endangered species. Rarity may emerge from secretive behaviors, small population sizes, or spatial clustering over large areas, such that encounters of the animal or evidence of its activity are infrequent relative to its probability of detection. Rarity poses many challenges in the estimation of abundance (W. Thompson 2013), including those related to inefficiencies in sampling. Small, highly clustered populations are difficult to sample, and may be costly when the organism of interest is absent from the majority of survey sites. Absence data are informative in analytical frameworks acknowledging imperfect detection such as capture-mark-recapture or occupancy frameworks. However, a dataset with a relatively large number of “zeroes” that arise from sampling in an area where the species is not present is costly because resources are spent gathering relatively poor information, and in addition it results in over-dispersed data that may not conform to standard statistical distributions, inhibiting inference (Barry and Welsh 2002; Martin et al. 2005). Notwithstanding an unbiased estimate, precision will be low in such situations, highlighting the importance of efficient sampling and analytical methods. Nevertheless, rare species represent an important concern for wildlife managers, usually because low population size implies a threat of extirpation or extinction, and rarity imparts a lack of critical information for management decisions.

Efficient sampling schemes for rare or patchily distributed species have been recently gaining in attention (W. Thompson 2013), but the problem has long been a focus of statistical ecology. Statistical treatments of the problem have been suggested through the use of zero-inflated Poisson or negative-binomial models (A. Welsh et al. 1996; Barry and Welsh 2002; Martin et al. 2005), wherein the abundance data are governed by a different distribution than the absence data, but others indicate that the separate modeling of absence data and abundance data implemented by the zero-inflated models has limited sensibility in terms of ecological or biological meaning (Kéry and Royle 2015). Alternatively, improvements to sampling design can attempt to address these difficulties. Stratified sampling can reduce variance in abundance estimates if the population can be partitioned by some well-defined homogeneous features; for example, if the organism is known to be strongly associated with a particular habitat type, estimator performance may be improved by allocating sampling proportionally to homogeneous habitat classes. Cluster or multistage sampling, and adaptive sampling take advantage of natural clustering of populations in order to improve inferences (S. K. Thompson 1990; Morrison et al. 2001; W. Thompson 2013).

Adaptive sampling strategies are a promising method for sampling small, clustered populations. Employing simple random sampling to estimate abundance of rare populations will result in zero-inflated data and estimators with poor precision (Salehi and Seber 1997). Often, the spatial structure of the population is not known or difficult to predict prior to surveys, making standard methods of stratification difficult to employ - adaptive sampling procedures operate only upon observed data and thus can be more effective than simple random or stratified sampling in these situations (S. K. Thompson 1990; Turk and Borkowski 2005). These methods result in preferential sampling in areas of higher animal abundance, and this framework can provide greater precision than simple random sampling among small, highly clustered populations (S. K. Thompson 1990; Salehi and Seber 1997; Brown 2003; Rapley and Welshy 2008; Smith et al. 2012). Certainly, preferential sampling – where the location of data collection is dependent upon the variable of interest – if unaccounted for can introduce bias into predictions made by the analysis. This has been examined generally by Diggle et al. (2010) and Pati et al. (2011), and recently incorporated into the ecological literature by Conn et al. (2017). Adaptive strategies are formulated to account for non-random sampling, either by using design-unbiased estimators under strict sampling protocols (Brown et al. 2013), or by modeling the selection of data locations jointly with the variable of interest (Conn, Thorson, and Johnson 2017).

The original design-based adaptive cluster sampling (S. K. Thompson 1990) begins by implementing an initial random sample of sites to survey - termed the primary sample. A critical threshold value of the state variable of interest is selected and secondary sampling sites are generated (or “triggered”) around each initial site wherever the threshold is met. Sampling resumes at each of these secondary sites, and additional sites can be triggered in turn if the threshold is met at these secondary sites. The chief advantage of this method is a distribution of effort that is a result of the spatial structure of the population. However, there exist drawbacks such as the randomness of the final sample size that make logistical constraints difficult to meet (Turk and Borkowski 2005), although there are methods that address this limitation (Christman and Lan 2001; Salehi and Seber 2002). There have been numerous contributions to variations of adaptive sampling allowing for flexible study design and inference under design-based estimators (Brown et al. 2013).

Recently, methods for studying patchily distributed populations have begun incorporating adaptive sampling principles into model-based sampling frameworks. A notable departure from the design-based adaptive sampling is the flexibility in modeling the primary sample. During the primary sample, an index variable is measured that is modeled to be conditional upon the state variable. Provided that the relationship between the index and state variables is satisfied, any valid statistical model may be used, including the ordinary case when the index variable is equivalent to the state variable. Selection of a threshold for adaptive sampling is based upon this index variable measured during the primary sampling occasion. For example, Pacifici et al. (2016; 2012) were concerned with enhancing occupancy model performance, and thus used occupancy models in the primary and secondary sampling occasions, selecting a threshold based upon detections out of K visits exceeding some *a priori* number. Conroy et al. (2008) used occupancy models in the primary sampling occasion, also selecting the threshold based upon detections exceeding some *a priori* number, and implemented capture-mark-recapture methodology in secondary sampling for estimation of vole abundance. Pacifici et al. (2016) made additional contributions to model-based adaptive sampling methodology by incorporating a spatially-explicit occupancy model along with the adaptive procedures, demonstrating a marked improvement in confidence interval coverage, particularly when sample sizes were sufficiently large to reliably estimate spatial covariance parameters.

Adaptive sampling is especially advantageous when the sampling protocol is time consuming or expensive regardless of whether or not individuals are detected. For example, in the use of technologies such as camera trapping or noninvasive genetic sampling, establishing sampling arrays or deploying highly trained dog teams to survey sample units is extremely costly, and this cost is incurred even if a sample size of 0 is obtained. As such, it is advantageous to integrate some sort of adaptive sampling strategy in such studies.

The combination of spatial capture-recapture and adaptive sampling is motivated by an effort to estimate the current population of moose (*Alces alces*) in the Adirondacks of northern New York, an area of approximately 24,000 km^2^. The moose population in this region is not growing according to management expectation, and the population was estimated at just 500-800 moose in the Adirondack park during 2010 (Wattles and DeStefano 2011). Spatial capture-recapture is an advantageous method to use for sampling moose in New York because it performs well with low-density, vagile species, and when coupled with non-invasive genetic identification of individuals from scat or hair, physically capturing individuals is unnecessary. A recent pilot study demonstrated the viability of using detection dogs to locate fecal pellets of moose in the Adirondack region (Kretser et al. 2016). A subsequent large-scale sampling effort was initiated across the Adirondacks of New York using random cluster sampling and detection of scat using trained dogs. Of 91 sites visited, 60 did not result in detection of moose scat throughout the sampling period; this was costly with respect to time, effort, and money. The geographic rarity of moose in this system warrants the evaluation of adaptive sampling as a method to enhance cost-efficiency and sample sizes, and improve statistical certainty in parameter estimates relevant to estimating the population size of moose.

In this paper, we develop an estimator of population density for spatial capture-recapture models under an adaptive sampling scheme in which the first stage of sampling produces an index to local population size and allows more intensive sampling to be triggered based on the observed index value. We investigate the performance of an adaptively sampled spatial capture-recapture model. We validate the model’s performance against a simulated population subjected to five sampling schemes, comparing the adaptive method to the ordinary SCR method when adaptive sampling is triggered by an index variable which is a function of site-specific abundance. We evaluate the performance of the adaptive sampling scheme by comparing it to an estimator based on conventional (non-adaptive) sampling based on a similar total sampling effort as well as estimators which ignore the index variable altogether. We also discuss the cost gains achieved by implementing an adaptive sampling framework and also potential application scenarios for which it might be useful.

We hypothesize that SCR implementation across all sites with index information will be unbiased with the smallest variance. Adaptively sampled SCR with index information is expected to be unbiased with greater variance than a full implementation of SCR at all sites, and the adaptively sampled SCR ignoring the index data (which results in biased sampling) is expected to be greatly biased. Under our expectations, we anticipate that the proposed method will be most useful in situations with highly patchy population distributions.

## Materials and Methods

### Spatial capture-recapture

Spatial capture-recapture extends ordinary capture-recapture models by formally integrating spatial information inherent in animal sampling data into the probability functions describing the detection process. Specifically, in its simplest form it makes the assumption that animal home ranges are distributed uniformly over the landscape and that detection of an individual is a function of distance from the home range centroid to the detector (Efford 2004). We describe the concepts of ordinary SCR, and then extend it to incorporate a model for the adaptive method.

### Data and sampling schemes

We simulate *g* ∈ {1, 2 *, …, G*} distinct populations or groups, each potentially sampled by a single transect (Figure 3); in so doing, referencing site or group is equivalent since the site samples one and only one population *g*. The transect is discretized into *j* ∈ {1, 2 *, …, J*} segments represented by the set of coordinates **x** at which animal encounters may occur. The variable *Ng* denotes the group-specific population size, and the total population is 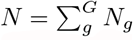. Any individual can belong to one and only one group, indexed for each individual by the vector 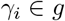, having length *N*.

**Figure 1:**
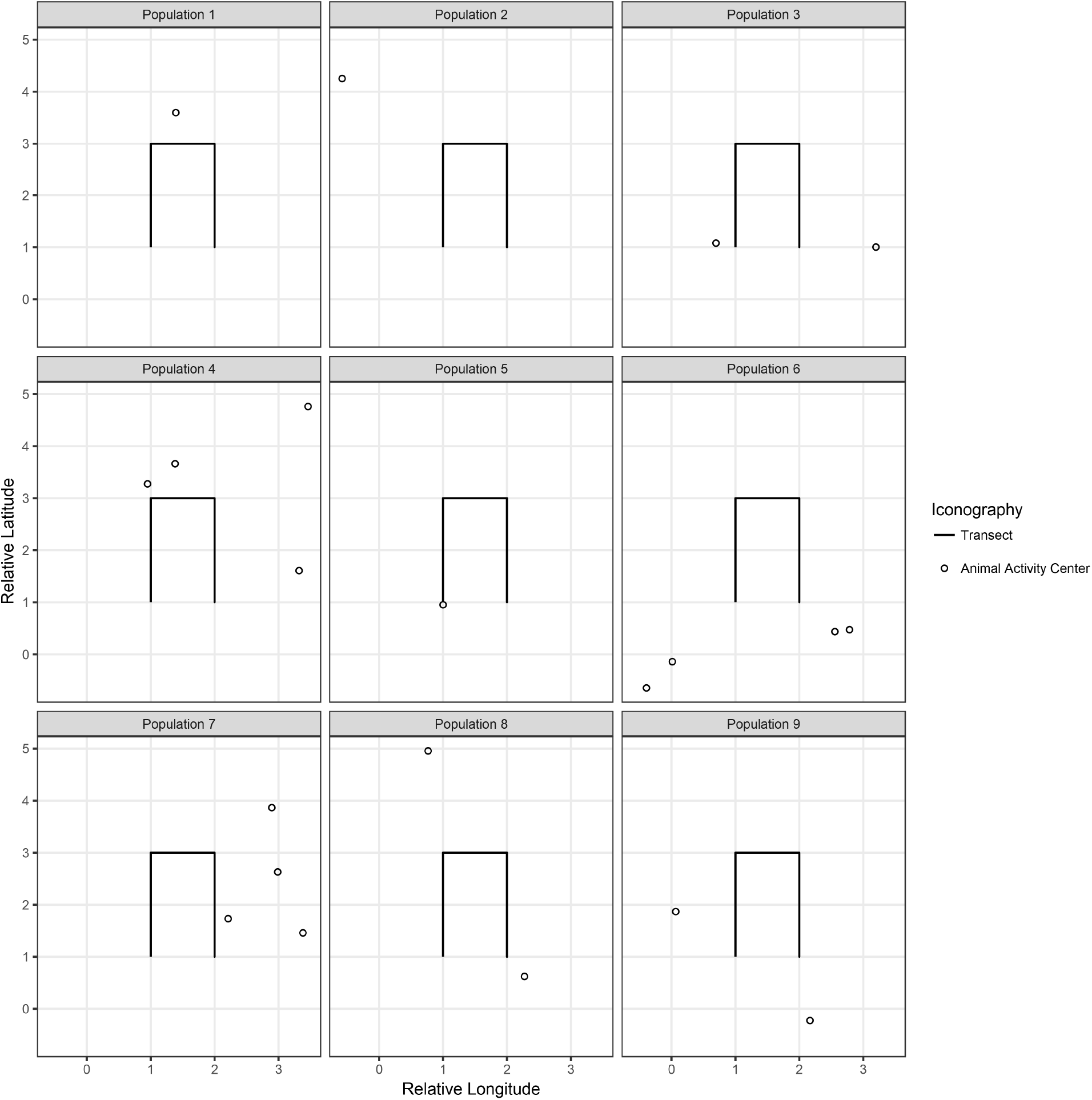
Independent animal populations are each sampled by a transect (solid line); 9 realizations of the simulation are displayed here. Animal activity centers (open circles) are distributed uniformly within the state space of each population. The population is simulated as a homogeneous Poisson point process with intensity equal to 2.

**Figure 2:**
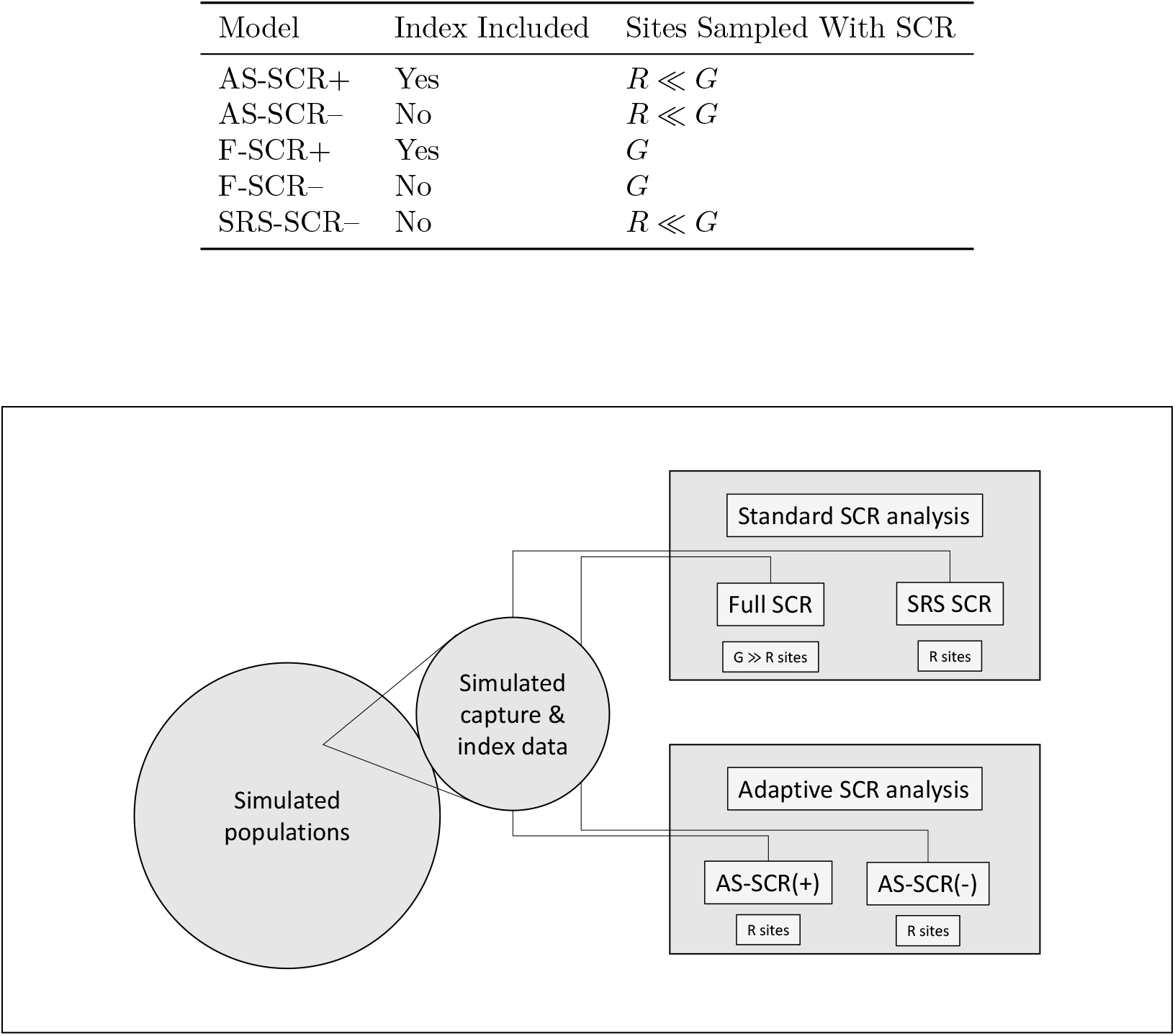
Four combinations of analytical and sampling methods are tested against simulated data. Two standard SCR analyses are performed on the full set of *G* sites, and also a subset of *R* randomly selected sites. Two adaptive SCR analyses are performed on *R* sites with the index exceeding a threshold *τ*. The plus and minus designations indicate whether index data is incorporated into the estimation procedure, or not, respectively.

**Figure 3:**
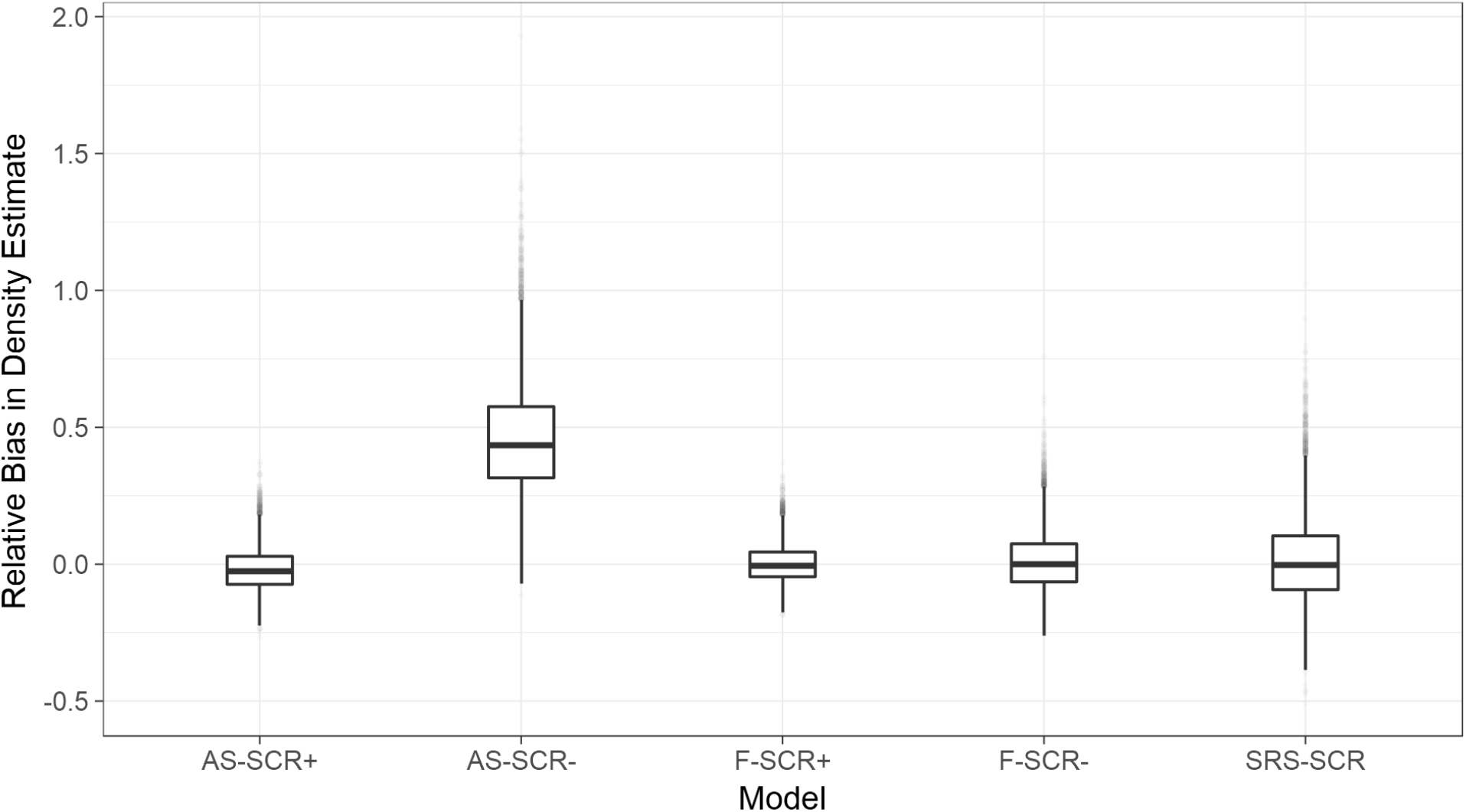
Graphical comparison of model performance. We show the distribution of relative bias in the density estimates for each model. The solid horizontal lines represent the median relative bias, the boxes represent the interquartile range, and outliers are represented by transparent dots.

For any particular group *g*, let *y_ijk_* denote a spatially-explicit encounter observation for individual *i* ∈ {1, 2 *, …, N_g_*} at trap or transect portion *j* during sampling occasion *k* ∈ {1, 2*, …, K*}. Accordingly, **y***_g_* denotes the matrix of encounter observations for each individual encounter at each traps within group *g*, also called the “encounter histories”. The group-specific structure is adopted from Royle and Converse (2014) and it assumes that each group population *N_g_* is mutually independent of one another. This is applicable in practice when trap arrays are spaced sufficiently far relative to the home range of the organism, and the sampling period is sufficiently short such that individuals do not occur in more than one group.

We propose a two-phase adaptive sampling spatial capture-recapture (AS-SCR) scheme; in the primary sampling phase, each of the *G* sites are sampled by some efficient method which produces an observation *τ_g_* which we assume is an index of local population size *N_g_*. The index could be a count obtained, for example, by road crossings of tracks of the species of interest, or counts of scat or other sign along roads or transects. The secondary phase samples a subset of size *R* of the *G* sites using a method that can produce individual animal encounters, with *R* ≤ *G*. The selection of this subset of sites is contingent upon whether the index *τ_g_* exceeds some prescribed threshold *T*: where *τ_g_* > *T*, the site *g* is sampled by capture-recapture to obtain individual animal encounters, and where *τ_g_* ≤ *T*, only the index observations are retained; the capture-recapture sampling is not implemented.

### Statistical models

#### SCR sites

At sites where *τ* > *T*, SCR is implemented, and we estimate the population size *N_scr_* as the sum of *N_g_* among these sites,

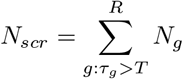

incorporating standard SCR techniques into the procedure. The point process model, detection model, and encounter model are described hereafter.

### Point process model

A key element of all SCR models is the introduction of a latent point process model which describes the distribution of individual home range centers or activity centers in the vicinity of the sampling array. We define *S_g_* to be the two-dimensional state-space of the activity centers **s**_*i*_ of all individuals *i* ∈ 1, …, *N_g_* within group *g*. We assume that the distribution of all **s***_i_* conditional on individual group membership 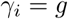 is

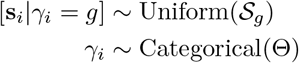

Where Θ is a vector of group-specific inclusion probabilities of length *G*. We use the notation [*Y |X*] to denote the probability mass function of a random variable *Y* conditional on *X*. This formulation is consistent with a Poisson point process, which may be extended using a linear combination of covariates representing attributes of the landscape. We operate under the assumption that the state space area is constant across all *g*. Additionally, take **S***_g_* to mean the set of individual activity center coordinates for all 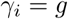.

The latent variable *N_g_* is well-defined for any explicit specification of the state-space *Sg*. Because all *G* populations are independent, each population has its own defined state space which is chosen to be a planar region containing each sampling array. The state space *Sg* of the latent activity center locations **s** should comprise the set of all possible coordinates that could have produced the data; however, integrating over an infinite array is not computationally practical, so the extent of the state-space is chosen large enough such that individuals with activity centers at the edge of the state space have a negligible probability of encounter, ensuring as near as possible that the expected frequency of encounter outside of the state space is 0 (J. A. Royle et al. 2014, 131–33).

### Detection model

The second key element of SCR models is the specification of a model for encounter probability of an individual with activity center **s** *i* near each trap or sampled unit. Most SCR models posit that the encounter probability *p* in a trap or transect portion *j* with known coordinate **x**_*j*_ is a function of distance between the individual’s activity center **s** *i* and the device. Sensible models for *pij* are monotonically decreasing whereby the value of *pij* decreases with increasing distance between **s**_*i*_ and **x**_*j*_. In our simulation study we use one of the most commonly used encounter probability models, based on the kernel of a Gaussian probability density function:

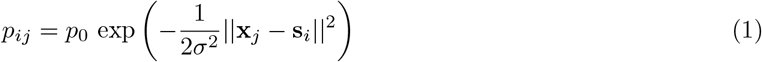

The term *p*_0_ denotes the baseline encounter probability when *||* **x**_*j*_ − **s**_*i*_ || = 0, and *σ* is a spatial scale parameter which relates probability of encounter of an individual in a trap to the distance between the trap *j* and **s***_i_*. This model serves to describe the probability of detection, which is one part of the encounter observation model described hereafter.

### Encounter observation model

Under a known-*N* scenario, the distribution of the encounter history data **y**_*g*_ is the product of *N_g_* × J_g_ *b* binomial probability mass functions:

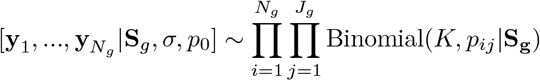

where *J_g_* represents the number of discrete transect portions within group *g*, and *p_ij_* is as it is defined in Equation 1.

At the time of the survey, *N_g_* and thus *N_scr_* are unknown, so we modify the structure of the model using the method of parameter-expanded data augmentation (J. A. Royle and Dorazio 2012). A super-population *M* is defined where *M* » *Nscr* and *M* ⊃ *Nscr*, and a new parameter *ψ* is included which describes the proportion of *M* that are part of the real population exposed to sampling. That is,

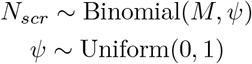

When the distribution for [*N_scr_|ψ*] is integrated over the prior for *ψ*, this is equivalent to establishing a marginal prior for the random variable *N_scr_* under a Bayesian mode of inference where *N_scr_~* Uniform(0, M). The formulation with the additional parameter *ψ* allows a more convenient representation of the encounter observation model through a zero-inflated binomial model. Ignoring group structure temporarily, the modified observation model is as follows:

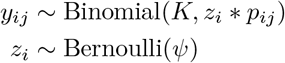

for *i* ∈ {1, 2, …, *M*} potential individuals. If the quantity *n* represents the number of observed individuals, this implies that an additional *M − n* all-zero encounter histories are added to the observation record with the goal of estimating what proportion of these *M − n* individuals are nonexistant or “structural zeros”, and which are true unobserved individuals or “sampling zeros”. The term *zi* denotes the inclusion index for each individual *i*, with *zi* = 0 for structural zeros and *z_i_* = 1 for sampling zeros. The super-population of *M* individuals serves as candidates to be included ( *z* = 1) in the population by the MCMC algorithm.

The likelihood for the individual observations is then

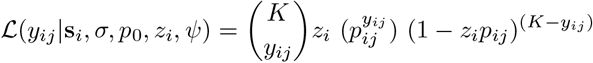

### Group membership and population model

To generalize this encounter model to a stratified population with *R* strata or groups, we represent an individual’s group membership as the variable 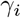 such that 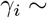 *~* Categorical(Θ). The term Θ is a vector of all group-specific probabilities *θ_g_ ∈ {θ*_1_, *θ*_2_, …, *θ_R_*}. Definition of the vector Θ is induced by the following population model:

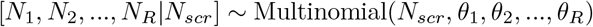

where 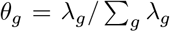. This is equivalent to the Poisson model for group-specific population sizes when conditioned on the total (among all groups) population size where the Poisson mean is

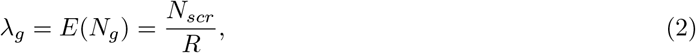

or the expected number of individuals within any group *g*.

To implement the data augmentation procedure, the *n* observed individuals are assigned to the group in which they were observed, and the *M − n* unobserved individuals are assigned to groups according to the group-specific probabilities in Θ, resulting in an augmented group-specific population size *M_g_* where *M_g_* > *N_g_*. Group-specific population size *Ng* does not appear explicitly in the likelihood; instead it is a derived parameter obtained by summing the values for *zi* for those individuals within group *g* such that

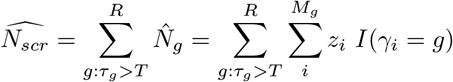

Where *I* is the indicator function evaluating to 1 if *γi* = *g* and 0 otherwise. We emphasize once again that the sites considered here are only those where *τ* exceeds the threshold *T*.

### Index observation model

For the model of index observations, the key assumption is that *τ_g_* must be conditional on *N_g_* in some fashion. In general, there is no explicit linkage of the index data *τ_g_* to *N_g_* of the SCR state-space. However, if the sample transect is the same between the primary and secondary sample, then in fact we can regard the index count as a thinning of the total encounter frequency of the SCR study. Other situations may yield satisfactory interpretations of the index; for example if the index sample is based on a transect randomly oriented through the state-space or more practically oriented in a way without first inspecting the state-space so as to avoid favorable or unfavorable portions of the state-space.

A sensible model for the index observations where *τ* > *T* (indeed, the classical “index assumption”) is consistent with the following model:

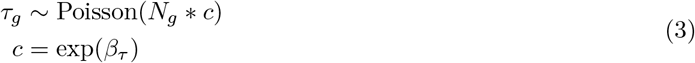

where *c* is some scalar adjustment parameter, and *β_τ_* is the underlying coefficient to be estimated. One might interpret the index variable as a thinning of the total encounter frequency of the SCR study, and the thinning rate is thus absorbed into *c*. In this manner, 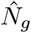 derived from the encounter observation model is used to inform the estimation of.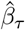

### Non-SCR sites

Notably, where *τ* ≤ *T* the second SCR sampling phase is not implemented, and so there are no encounter observations yielding information about *N_g_* at those sites; a distinct model must be used to estimate *N_g_*. The model for these sites relies on the propagation of information about *N_g_* from the expected population size *λ_g_* as well as from the index observations with the following relationship:

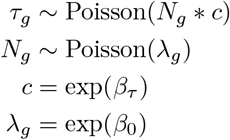

The expected population size is indirectly informed from the encounter observation model through *θg*, which is in turn informed by the observed individual group membership 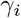. Other natural index models are possible, which we discuss later in the discussion.

For the total population size at all *G − R* of the sub-threshold sites, we indicate:

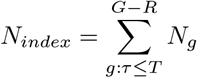

and the total number of individuals estimated at all *G* sites is:

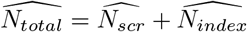

### Joint distribution of the data

The joint distribution of the index data and the SCR data is represented by the following:

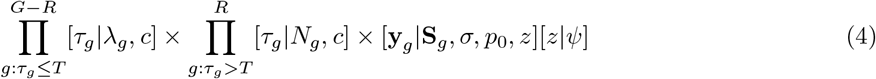

The distribution of the index data at the sub-threshold sites appear as the left term, and the distribution of the index data and the encounter data appear as the right term.

For sampling schemes ignoring the index observations, the terms containing *τ_g_* are removed from the joint distribution, and what remains is the typical observation model for group-specific SCR:

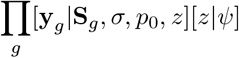

### Prior distributions

We have established one prior distribution previously – that of 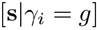, being assumed to be uniform over the state space of *S_g_*. Here, we establish prior distributions for the remaining variables.

The baseline detection probability parameter has prior distribution *p*_0_ *~* Uniform(0, 1) (assuming geographic coordinates are scaled to units of km).

The spatial scale parameter has prior distribution *σ* ~ Uniform(0, 10).

The data augmentation parameter has prior distribution *ψ* ~ Uniform(0, 1).

The mean group-specific population size has prior distribution *λ_g_* = exp(*β*_0_) and *β*_0_ *~* Normal(0,0.01).

The scalar multiplier has prior distribution *c* = exp(*β_τ_*) and *β_τ_~* Normal(0,0.01).

### Joint posterior distribution

The joint posterior distribution of the model parameters conditional on the observed encounter history data and index data is the product of the joint distribution for the data, and the prior distributions:

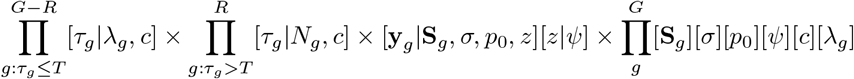

For the cases where adaptive sampling is not done, so that we only have SCR encounter history data, the terms including *τ* are removed, leaving the posterior distribution for a basic SCR model:

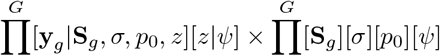

## Simulation Study

### Design

We evaluate the situation where *λ_g_* = 2 for all *g*, specified by the relationship in Eq.(2). This parameterization appears reasonable to the authors for studies of animals that occur at relatively low densities. We use the threshold value *T* = 4, and we simulate the index values *τ_g_* according to the relationship in Eq.(3), where *N_g_* is the result of the simulated group population size, and *c* = 3. For each simulation we generated *G* = 100 populations sampled on *K* = 3 occasions with a single rectangular transect represented as point detectors spaced at 0.1 unit intervals along its length (Figure 1). A total of 1000 simulations were generated with these parameter settings.

For added generality, the reader may use a gamma-Poisson mixture to model populations more dispersed than the Poisson scenario – the description of this model formulation can be read in Royle et al. (2012).

### Model configurations

Five model analyses are used to evaluate three sampling schemes (AS-SCR, F-SCR, and SRS-SCR, defined below) that either consider or ignore the index measurements obtained in the primary sampling phase. We evaluate the sampling schemes of AS-SCR considering (*AS-SCR+*) and ignoring (*AS-SCR–*) index observations, F-SCR considering (*F-SCR+*) and ignoring (*F-SCR–*) index observations, and SRS-SCR without index observations (*SRS-SCR–*).

*AS-SCR+* is the proposed adaptive sampling procedure that we test for bias and precision. We expect to observe unbiased parameters estimates, particularly estimated population size.

*AS-SCR–* represents an preferential sampling situation and we expect it should be positively biased for density, because it samples only high-density areas without integrating information obtained from indices that fall below the index threshold.

*F-SCR+* is the ideal and most costly sampling procedure, incorporating SCR and index observations at every site, and it represents the most accurate and precise baseline to which we compare *AS-SCR+*.

*F-SCR–* represents a standard SCR procedure implemented at all sites without the index data, which we expect will resemble *F-SCR+* with a slight loss in precision. This is an ordinary application of SCR against which we test our new procedure.

*SRS-SCR–* represents application of standard SCR at a simple random sample of sites equal to the corresponding number of sites sampled by AS-SCR. It makes no use of index data. This comparison is important to include because it is a sampling method which a cost-restrained survey could adopt having little initial information regarding the local population. We expect this will be unbiased, but also that it will have the poorest precision in comparison to the other models in the set due to the relatively sparse information it gathers.

For each analytical procedure, we calculated relative bias, mean squared error (MSE), and 95% coverage for the parameter estimates. We transformed estimates of abundance into density to make them comparable across all models and sampling procedures. We report these summary statistics for estimates of density 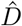, scale parameter 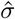, and baseline detection probability 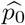, the three main parameters of interest in SCR.

Relative bias was calculated as:

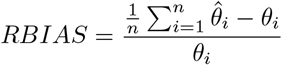

Where 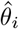 is the posterior estimate for parameter *θ* at simulation *i* with true value *θ_i_*.

MSE was calculated as:

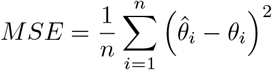

For reporting coverage, we estimated standard error using the Monte Carlo standard error estimator, used to construct Bayesian 95% credible intervals.

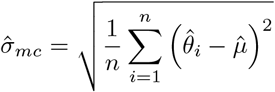

Where, 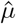 is the sample mean of all posterior estimates for parameter *θ* such that 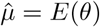 assuming the stationary distribution is achieved at all simulations.

### Bayesian analysis

We evaluate these models numerically with the JAGS software implemented in R with the package “jagsUI”. Ten-thousand MCMC iterations were performed for each model analysis with burn-in ranging from 500-1000 iterations. The software sampler was allowed to adapt under default settings.

## Results

As anticipated, percent bias and MSE are positive and high for the preferential sampling procedure that disregards the index data (AS-SCR–), with abundance overpredicted by approximately 46% on average (Table 1). Additionally, coverage is very low, with the 95% credible intervals for 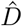 intersecting the true value 2.1% of the time (Table 2). Values of 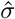 and 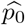 are unaffected by the omission of index information.

**Table 1:** Relative Bias for Parameter Estimates

| Model | $\hat{D}$ | $\hat{\sigma}$ | $\hat{p}_0$ |
| --- | --- | --- | --- |
| AS-SCR+ | -0.0198 | 0.0284 | 0.0056 |
| AS-SCR- | 0.4586 | 0.0045 | -0.0021 |
| F-SCR+ | 0.0025 | 0.0091 | -0.0003 |
| F-SCR- | 0.0128 | 0.0045 | -0.0008 |
| SRS-SCR | 0.0183 | 0.0077 | 0.0023 |

**Table 2:** Mean Squared Error for Parameter Estimates

| Model | $\hat{D}$ | $\hat{\sigma}$ | $\hat{p}_0$ |
| --- | --- | --- | --- |
| AS-SCR+ | 0.0000 | 0.0071 | 0.0001 |
| AS-SCR- | 0.0011 | 0.0093 | 0.0001 |
| F-SCR+ | 0.0000 | 0.0048 | 0.0001 |
| F-SCR- | 0.0001 | 0.0072 | 0.0001 |
| SRS-SCR | 0.0001 | 0.0128 | 0.0002 |

**Table 3:**
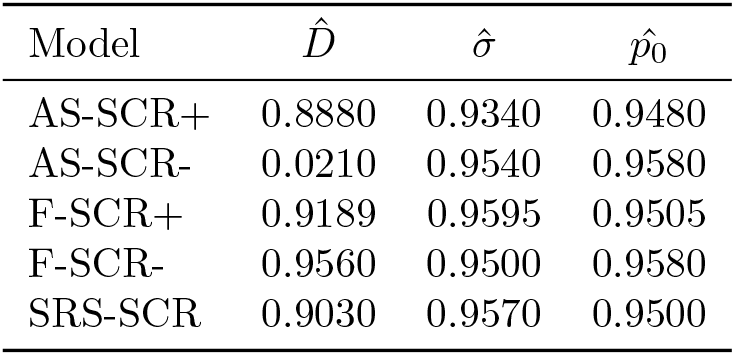
Coverage for Parameter Estimates over 95% credible interval

In comparison, our new adaptive sampling method (AS-SCR+) has low bias and MSE; bias is within 2% of the true density value, and coverage is high, with the true value included within the 95% credible interval approximately 89% of the time.

The results for the baseline comparisons (F-SCR+; F-SCR–; SRS-SCR–) are unsurprising; their estimates for all parameters are unbiased and have very high coverage. Notably, the bias, MSE, and coverage for 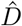 are nearly equal between AS-SCR+ and SRS-SCR–, but the standard deviation in the estimate under AS-SCR+ is nearly half that under SRS-SCR– (Table 4). As F-SCR+ is the most information-rich, it is unsurprising that its bias, MSE, and estimate standard deviations are the smallest.

**Table 4:**
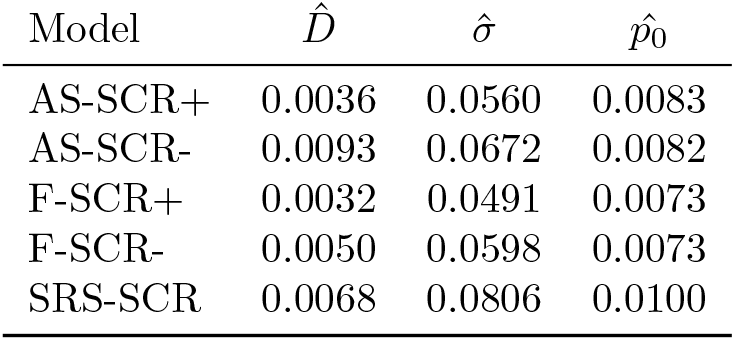
Monte Carlo Standard Error for Parameter Estimates

Under the Poisson random population simulation setting, the adaptive procedure sampled one-half of the total number of sites compared to the full treatment with virtually no loss in estimation accuracy or precision. The AS-SCR+ estimator also estimated density with a higher precision than the equal-sized SRS-SCR application.

## Discussion

### Discussion Overview

We present an adaptive sampling approach to spatial capture-recapture sampling that can be used to increase sample sizes for a nominal cost or reduce the cost in obtaining a target sample size, as compared to a simple-random-sample application. This application is especially relevant for low-density, patchily distributed species, which typically present significant logistical and statistical challenges.

The adaptive method distributes sampling effort in accordance to the spatial structure of the population, ideally minimizing sampling in areas of low population density. Pacifici et al. (2016) demonstrate that incorporating a spatially-explicit model for the variable of interest enhances parameter estimation, either by spatial process models or by spatial random effects (also explored by Johnson et al. (2013)). The incorporation of spatial capture-recapture with adaptive sampling may be preferable to a non-spatial model. The formulation of spatial capture-recapture in this simulation study used a homogeneous Poisson point process to represent the animal activity centers, but it could be easily extended to incorporate an inhomogeneous Poisson point process in which the assumption of uniform density is relieved, potentially allowing for more refined prediction of abundance to unsampled areas (J. A. Royle et al. 2014).

Our results suggest that adaptive SCR is equally effective as a full treatment of SCR under a Poisson randomly distributed population. While we did not perform a formal cost analysis, it is easy to consider the potential cost-savings of reducing the number of distinct sites to visit. In the motivating context, the daily cost incurred by scat detection dog surveys was approximately $1000 USD daily in 2016, whether or not moose were present on the transect. Transects with no scats composed approximately 2/3 of all the transects visited, and 91% of these remained without scat throughout the sample period. At maximum efficiency four sites could be visited per day, making the per-site cost at least $250; with 40 unoccupied sites, the cost of allocating effort to these sites was at least $10,000, or 10% of the summer’s expenditures. Under an ordinary SCR survey, equal cost and effort is spent at unoccupied sites as occupied sites, reducing efficiency when unoccupied sites compose the majority of sites visited. After implementing the adaptive SCR procedure in 2017, we observed a four-fold increase in moose fecal sample collection with no additional effort, indicating the substantial benefits in applying the method in cases where the organism of interest is sparse on the landscape.

Previous studies of design-based adaptive cluster sampling methods are known to be sensitive to the choice of adaptive threshold (Turk and Borkowski 2005; Brown 2003). The selection of the threshold value directly affects the resulting within- and between-network variances, and due diligence is required to maximize efficiency of the adaptive procedure over a simple-random-sample implementation. A similar application of adaptive sampling to model-based sampling frameworks by Conroy et al. (2008) indicates relative insensitivity of abundance estimates to a range of threshold values. However, the flexible structure of our framework precludes a general suggestion; selection of a proper index value is best informed by pilot surveys and simulation under the most applicable index model for the study system evaluated on a case-by-case basis.

The specific type of index should be carefully selected such that there is some quantifiable relationship between the index and local population density. For example, in the motivating context, the index variable was the number of scat piles encountered on the first visit to the site. Selection of a poor index measure will not bias the spatial capture-recapture parameter estimates, but sampling conditional upon this index measure – if truly uncorrelated with the population – will likely result in a sampling procedure resembling a simple random sample without an *a priori* defined number of sites visited.

### Extension of Results

The adaptive SCR model can accept many varieties of index model. We considered our model suitable to describe fecal deposition as a function of population size. Alternative models can vary to accommodate the study system. For instance, one may wish to formulate the model conditional on detection/non-detection of an animal:

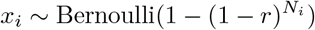

Where, *x_i_* is the detection of a species at site *i*, *r* is an individual detection rate, and *N_i_* is the local population size. One might also wish to integrate occupancy as the index model as done in Conroy et al. (2008) or Pacifici et al. (2016) and select a threshold of the occupancy estimate or raw initial count to trigger adaptive SCR.

Theoretically, the index model could also be based upon citizen science data, increasing the applicability of this model at wide spatial extents and further reducing the cost burden on researchers. For example in New York, occupancy of moose was estimated using reported observations from hunters (Crum et al. 2017), and could be augmented through the use of citizen science apps such as iSeeMammals (Sun et al. iseemammals.org) which collects presence/absence observations of mammals from hikers and camera stations. The adaptive site selection could proceed by selecting a threshold of estimated occupancy across the survey area, reducing the burden of index data collection to the researchers. The potential for sampling or detection bias in citizen science is recognized (Kéry et al. 2010), so we expect that some collection of index data would still be required from the researchers to minimize this bias, but further research in this area is warranted.

Application of AS-SCR may be particularly useful when covariates affecting density are weakly correlated or unknown, permitting efficient investigation of novel study systems since the resulting distribution of sampling effort is conditional upon observed data. Alternatively, if covariates are known that are strongly correlated with density, it may be possible to condition the index model on these, eliminating the need to conduct a preliminary index survey. This possibility also warrants further testing.

Spatial capture-recapture has been demonstrated to be more effective than non-spatial methods in estimating density of rare and elusive organisms, such as large carnivores, than non-spatial methods (Kéry et al. 2011; Sollmann et al. 2013; Blanc et al. 2013). However, data requirements for spatial capture-recapture are larger than ordinary capture-recapture methods owing to the additional parameters to estimate, so there is a larger trade-off between sampling intensity and the spatial extent that may be surveyed. This problem is exacerbated by low-density populations, where data richness is not even across the survey area. The adaptive method actively achieves focused sampling intensity at sites with greater data richness, allowing for greater potential precision than a simple random sample could typically achieve under the same circumstances. We suggest that the application of the adaptively sampled SCR method can be particularly useful when it is critical to reduce cost and effort to meet budget or time constraints while at the same time maintaining reliable parameter estimates.

## Acknowledgements

We would like to acknowledge the funding and guidance provided by the New York State Department of Environmental Conservation, Conservation Canines for their assistance in collecting the moose scat data, and the Wildlife Conservation Society for assistance in landowner access and field logistics.

Any use of trade, firm, or product names is for descriptive purposes only and does not imply endorsement by the U.S. Government.

